# PAT2 regulates autophagy through vATPase assembly and lysosomal acidification in brown adipocytes

**DOI:** 10.1101/2020.01.24.918078

**Authors:** Jiefu Wang, Martin Krueger, Stefanie M. Hauck, Siegfried Ussar

## Abstract

Brown adipose tissue (BAT) plays a key role in maintaining body temperature as well as glucose and lipid homeostasis by its ability to dissipate energy through mitochondrial uncoupling. To facilitate these tasks BAT needs to adopt its thermogenic activity and substrate utilization to changes in nutrient availability, regulated by a complex network of neuronal, endocrine and nutritional inputs. Amongst this multitude of factors influencing BAT activity changes in the autophagic response of brown adipocytes are an important regulator of its thermogenic capacity and activity. Increasing evidence supports an important role of amino acid transporters in mTORC1 activation and the regulation of autophagy. However, a specific role of amino acid transporters in BAT regulating its function has not been described. Here we show that the brown adipocyte specific proton coupled amino acid transporter PAT2 rapidly translocates from the plasma membrane to the lysosome in response to amino acid withdrawal, where it facilitates the assembly of the lysosomal vATPase. Loss or overexpression of PAT2 therefore impair lysosomal acidification, autophagolysosome formation and starvation induced mTORC1 activation.

## Introduction

Brown adipose tissue (BAT), with its unique ability to dissipate excessive energy in form of heat through mitochondrial uncoupling, plays an important role in regulating body temperature, but also glucose and lipid homeostasis and consequently body weight (Klepac, Georgiadi et al., 2019, Nedergaard & Cannon, 2018, Townsend & Tseng, 2014). Importantly, BAT activity itself is tightly regulated by environmental signals, as well as the metabolic state of the organism (Hankir & Klingenspor, 2018, Heeren & Scheja, 2018, Hoeke, Kooijman et al., 2016, Li, Schnabl et al., 2018, Mills, Pierce et al., 2018, Oelkrug, Polymeropoulos et al., 2015, Okla, Kim et al., 2017, Ramirez, Lynes et al., 2017). This underscores the important role of BAT as rheostat sensing the organismal state to regulate whole body metabolic function through a complex network of neuronal, endocrine and nutritional inputs. The role of the sympathetic nervous system, as well as glucose, fatty acids and other metabolites have been extensively described in the regulation of BAT activity (Hankir, Cowley et al., 2016, Hankir & Klingenspor, 2018, Heeren & Scheja, 2018, Hoeke et al., 2016, Kuruvilla, 2019). However, surprisingly little is known about the potential role of amino acids in the regulation of BAT function. *A*lanine was shown to inhibit glucose oxidation of brown adipocytes (Lopez-Soriano & Alemany, 1989), whereas leucine as well as arginine appear to promote BAT growth and function (Wanders, Stone et al., 2015, Wu, Satterfield et al., 2012).

Cellular amino acid levels are sensed and regulated by a complex network of proteins and organelles centered around mTORC1 activity (Condon & Sabatini, 2019). Conditional ablation of raptor in adipocytes resulted in increased lipolysis and lipophagy, which could be rescued by inhibition of autophagy through depletion of ATG7 (Zhang, Wu et al., 2019). Autophagy is a general degradation process through the delivery of various intracellular structures to the lysosome for degradation in response to cellular stress (Dikic & Elazar, 2018, Galluzzi, Pietrocola et al., 2014, Mizushima, 2018), whereby the proteolytic activity of the lysosome itself depends on vATPase mediated luminal acidification (Kissing, Hermsen et al., 2015). Upon hydrolysis, amino acids are released from the lysosome into the cytoplasm where they activate lysosomally targeted mTORC1 (Yu, McPhee et al., 2010) to regulate a multitude of cellular processes (Saxton & Sabatini, 2017). In this context, increasing evidence highlights the importance of lysosomal amino acid transporters in mTORC1 activation and the regulation of autophagy (Broer & Broer, 2017, Goberdhan, Wilson et al., 2016, Rebsamen, Pochini et al., 2015, Wyant, Abu-Remaileh et al., 2017). Autophagy regulates adipocyte differentiation and thermogenic gene expression (Ferhat, Funai et al., 2018). However, a specific role of amino acid transporters in BAT regulating its function has not been described.

We previously identified the proton coupled amino acid transporter PAT2 (SLC36A2) as highly enriched in brown adipocytes (Ussar, Lee et al., 2014). PAT2 is a proton coupled amino acid transporter that belongs to the SLC36 family (Schioth, Roshanbin et al., 2013, Thwaites & Anderson, 2011), with very narrow substrate specificity (Rubio-Aliaga, Boll et al., 2004) and strong pH dependence (Boll, Foltz et al., 2002, Foltz, Oechsler et al., 2004, Kennedy, Gatfield et al., 2005, Rubio-Aliaga et al., 2004). We showed that in contrast to PAT1, PAT2 does not localize to the lysosome, but is found at the plasma membrane of fully differentiated brown adipocytes (Ussar et al., 2014). However, the function of PAT2 in brown adipocytes is not known. Here we show that PAT2 resides at the cell surface of mature brown adipocytes to sense extracellular amino acid levels, as depletion of extracellular amino acids results in rapid translocation of PAT2 form the cell surface to the lysosome. We show, that PAT2 at the lysosome interacts with the V0 subunit of the vATPase facilitating full assembly of the enzyme by recruiting the cytosolic V1 subunit, as well as regulating pumping efficiency of the vATPase. Deregulation of PAT2 by either overexpression of knockdown result in hyper- or hypoacidification of the lysosome, respectively, with profound effects on autophagolysosome formation and activation of mTORC1.

## Results and Discussion

To study the fasting response of metabolically important tissues, 8 weeks old chow diet fed male wildtype C57Bl/6 mice were fasted overnight. Overnight starvation significantly reduced blood glucose levels but did not impair body weight or weights of individual tissues (**Fig. S1A**).

Fasting did not change the expression of LC3b in BAT, or any other tissue investigated. In contrast, expression of the brown/ beige adipocyte specific genes uncoupling protein-1 (UCP-1) and amino acid transporter PAT2 (slc36a2) was significantly reduced (**Fig. S1B**). Albeit BAT showed no change in the expression of LC3b, fasting resulted in a strong conversion from the cytosolic LC3 type I to the autophagosome incorporated LC3 type II in skeletal muscle (tibialis anterior; TA) and brown adipose tissue (BAT), whereas liver upregulated LC3 level in general, but did not show increased conversion from LC type I to type II. These changes were not observed in subcutaneous (SCF) and perigonadal fat (PGF) (**Fig. S1C**). The increase in LC3 type II in BAT but not WAT was also confirmed by immunofluorescence stainings of LC3 (**Fig. S1D**), indicating that BAT is as sensitive to starvation as skeletal muscle. However, UCP-1 protein levels, in contrast to mRNA levels, were not reduced, but even appeared increased following an overnight fast (**Fig. S1C**), suggesting a complex role of starvation in mitochondrial uncoupling and function. We previously identified PAT2 as highly expressed in brown and beige adipocytes (Ussar et al., 2014) and the co-regulation with UCP-1 in response to an overnight fast prompted us to investigate its potential role in orchestrating the amino acid related fasting response. To study the function of PAT2 in brown adipocytes, we established brown preadipocyte cell lines stably overexpressing HA-tagged PAT2 (PAT2-HA) or depleted of PAT2 (shPAT2) (**Fig. S2A**).

As previously reported (Ussar et al., 2014), PAT2 expression is very low in preadipocytes and strongly induced upon brown adipocyte differentiation (**Fig. S2A**). Interestingly, protein levels of stably overexpressed PAT2 were also much lower in brown preadipocytes compared to fully differentiated mature brown adipocytes (**Fig. 1A**). Lysosomal protein degradation is the default pathway for protein turnover of cell surface proteins and plays an important role in regulating mTORC1 activation and autophagy (Abu-Remaileh, Wyant et al., 2017, Perera & Zoncu, 2016, Wu, Zhao et al., 2016, Zoncu, Bar-Peled et al., 2011). Indeed, PAT2 predominantly localized to lysosomes in preadipocytes (**Fig. 1B**). Thus, we tested if the observed differences in PAT2 protein levels between preadipocytes and adipocytes are the result of increased lysosomal protein turnover in preadipocytes. Treatment of brown preadipocytes with the vATPase inhibitor bafilomycin A1, preventing lysosomal acidification, increased PAT2-HA protein levels in preadipocytes (**Fig. 1C**) and resulted in accumulation of PAT2-HA in late endosome like multivesicular structures (**Fig. 1D**). Furthermore, insulin and the mTORC1 inhibitor rapamycin also, although to a lesser extent, increased PAT2-HA protein levels (**Fig. 1C** and **D**). A combination of bafilomycin and rapamycin with or without insulin resulted in detectable cell surface localization of PAT2 in preadipocytes (**Fig. S2B**). In contrast, stimulation with the β3 adrenergic receptor agonist CL316243 showed no effect compared to control cells (**Fig. 1C**). Together, these results indicate that the protein levels and subcellular localization of PAT2 are tightly connected to the intracellular amino acid sensing machinery. Furthermore, we observed increased proliferation in PAT2-HA compared to Scr and shPAT2 preadipocytes (**Fig. 1E**), suggesting also a functional connection between PAT2 and mTORC1 activity (Ben-Sahra & Manning, 2017). Indeed, previous reports have suggested a role of PAT2 in the regulation of mTORC1 (Suryawan, Nguyen et al., 2013), albeit no mechanistic details have been reported until now. Co-immunoprecipitation experiments revealed an interaction of PAT2-HA with mTOR and RagC, both components of mTORC1 (**Fig. 1F**) suggesting that PAT2 could directly regulate mTORC1 activity in preadipocytes.

**Figure 1:**
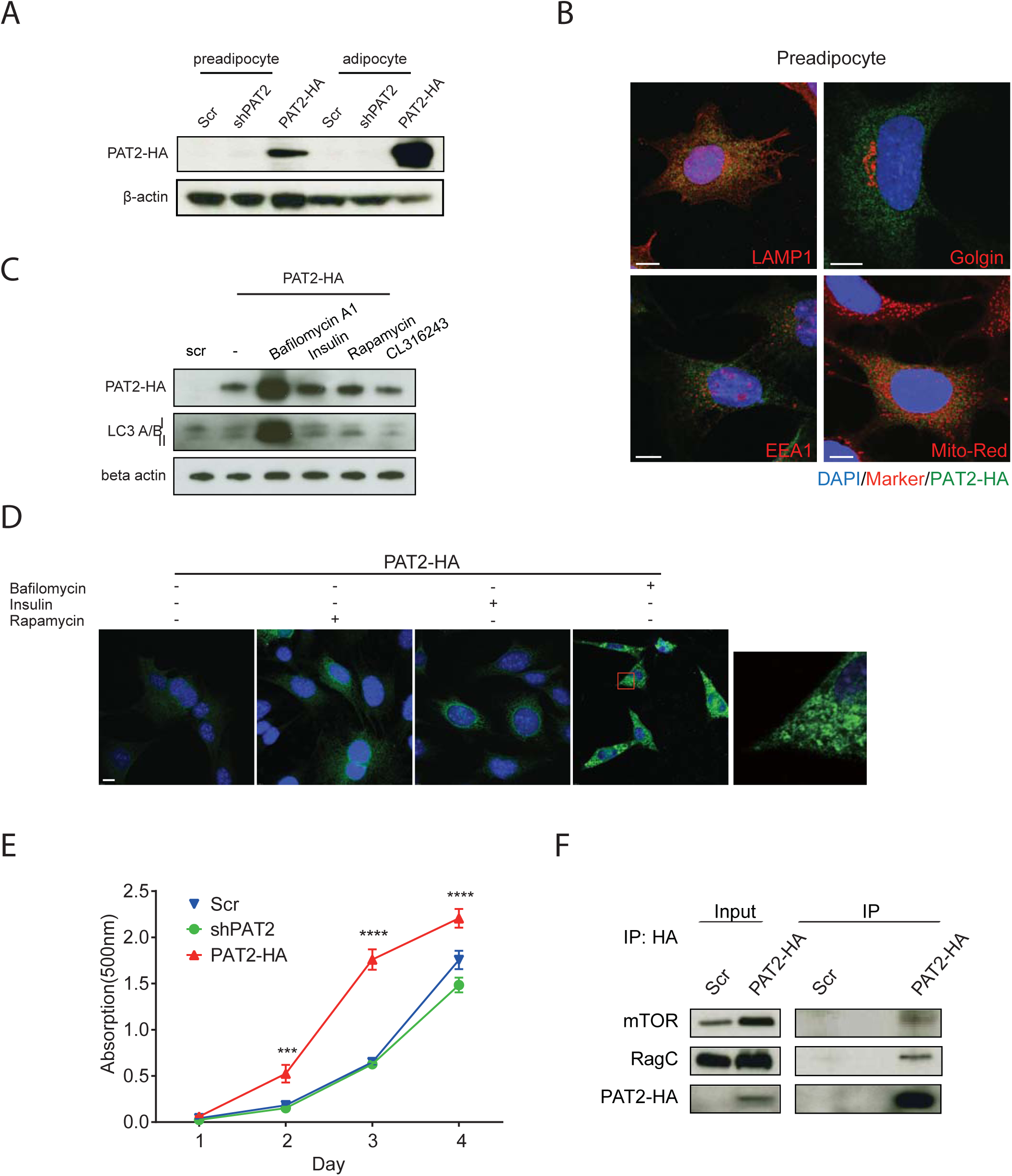
PAT2 is rapidly degraded in the lysosome in preadipocytes. **(A)** Western blot for PAT2-HA at day 0 (preadipocytes) and day 8 (brown adipocytes) of differentiation. **(B)** Immunofluorescence staining of PAT2-HA and different organelle markers (LAMP1 for lysosome; Golgin for golgi; EEA1 for late endosome, Mito-Red for mitochondria) in brown preadipocytes. Scale bar shows 7.5 µm. **(C)** Western blot for Scr and PAT2-HA brown preadipocytes treated with bafilomycin A1 (100nM), insulin (100nM), Rapamycin (10µM) or CL316243 (500nM). **(D)** Immunostaining for PAT2-HA in preadipocytes treated with Bafilomycin A1 (100nM), insulin (100nM), Rapamycin (10µM). Scale bar shows 10 µm. **(E)** Preadipocyte proliferation for Scr, shPAT2 and PAT2-HA brown preadipocytes. n=3. *** p<0.001, **** p<0.0001, RM two-way ANOVA with Tukeys’ post hoc test; error bars show SEM. **(F)** Co-immunoprecipitation of PAT2-HA with components of mTORC1 in preadipocytes.

However, endogenous levels of PAT2 are very low in preadipocytes and most of the protein appears to be readily degraded in the lysosome. In contrast to this, PAT2 predominantly localizes to the cell membrane in mature brown adipocytes (**Fig. 2A**) (Ussar et al., 2014). To understand a possible role of PAT2 in mature adipocytes in the regulation of mTORC1 activity and lysosomal function we differentiated PAT2-HA, shPAT2 and control cell lines. Knockdown or overexpression of PAT2 did not impair lipid accumulation (**Fig. S3A and B**) and expression of the key adipogenic transcription factor PPARγ (**Fig. 2B** and **C**), despite increased mRNA levels of PPARγ at day 8 in PAT2-HA brown adipocytes. In contrast, mRNA and protein levels of the brown adipocyte specific protein UCP-1 were elevated in both PAT2-HA and shPAT2 adipocytes compared to control cells (**Fig. 2 B** and **C**).

**Figure 2:**
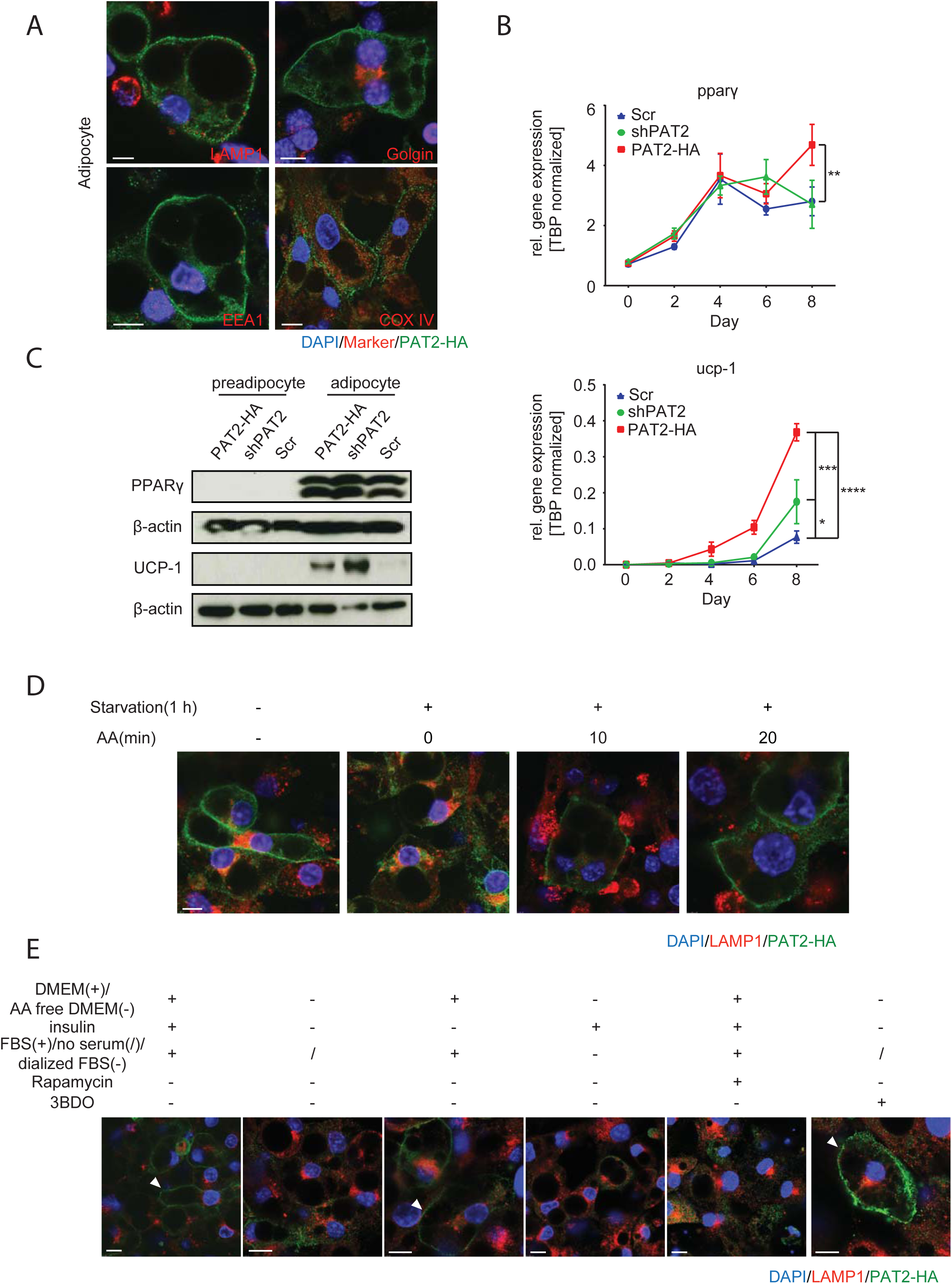
Subcellular localization of PAT2 in brown adipocytes depends on mTORC1 activity. **(A)** Immunofluorescence staining of PAT2-HA and different organelle markers (LAMP1 for lysosome; Golgin for golgi; EEA1 for late endosome, COX IV for mitochondria) in brown adipocytes (day 8). Scale bar shows 7.5 µm. PPARγ and UCP-1 mRNA **(B)** and protein **(C)** levels during differentiation (n=3-6 for Semiquantitative PCR; preadipocytes: day0; adipocytes day 8). **(D)** Immunostaining of PAT2-HA and LAMP1 in PAT2-HA brown adipocytes under normal culture conditions or amino acid and serum starvation for one hour with or without restimulation with amino acids for 10 or 20 minutes. Scale bars show 7.5 µm. **(E)** Immunostaining of PAT2-HA and LAMP1 in PAT2-HA brown adipocytes treated with 10 µM rapamycin, 100 nM insulin, 60 µM 3BDO for one hour. Arrows indicate plasma membrane staining. Scale bars show 10 µm.

As stated above, PAT2, as previously reported (Ussar et al., 2014), predominantly localizes to the cell surface in mature brown adipocytes (**Fig. 2A**). However, we found that serum and amino acid depletion for one hour induced a translocation of PAT2 from the cell membrane to the lysosome, which was rapidly reverted by stimulation with amino acids (**Fig. 2D**). The selectivity of the observed translocation to amino acid depletion was confirmed by inducing translocation using amino acid withdrawal in the presence of 10% dialyzed FBS (**Fig. 2E**). Moreover, the plasma membrane localization of PAT2 appeared to be dependent on mTORC1 activity, as rapamycin, similar to amino acid withdrawal, triggered internalization and lysosomal translocation, whereas activation using the mTORC1 activator 3BDO maintained membrane localization upon amino acid deprivation. In contrast, removal of insulin did not trigger the translocation of PAT2 form the cell membrane to the lysosome (**Fig. 2E**).

The data above, clearly establish the dependency of PAT2 localization and lysosomal degradation on mTORC1 activity, but also suggest a role of PAT2 in regulating mTORC1 function. Prolonged amino acid depletion resulted in rapid dephosphorylation and autophagy dependent rephosphorylation of S6K in control adipocytes (Yu et al., 2010). In contrast, prolonged starvation resulted in impaired reactivation of S6K in both shPAT2 and PAT2-HA brown adipocytes (**Fig. 3A** and **B**), while phosphorylation of mTOR was not changed (**Fig. S4A**). The difference between S6K and mTOR phosphorylation suggests that loss or gain of PAT2 alters autophagolysosomal amino acid release rather than growth factor signaling. Electron microscopy did not indicate differences in autophagosome numbers between the three cell lines (**Fig. 3C** and **D**), indicating that overexpression or depletion of PAT2 does not impair autophagosome formation. In line with this, we also did not observe differences in LC3 gene expression upon overexpression or depletion of PAT2 (**Fig S4B**). However, differences in mean autophagosome size were observed upon starvation (**Fig. S4C**). Thus, we tested if PAT2 regulates autophagosome and lysosome fusion. Co-immunostainings of LAMP1 and LC3 revealed reduced co-localization of these lysosome and autophagosome markers, respectively, in shPAT2 and PAT2-HA cells (**Fig. 3E** **and Fig. S4D**). Similarly, measuring proton dependent EGFP quenching in the autophagolysosome using a transiently transfected mCherry-EGFP-LC3 construct (N’Diaye, Kajihara et al., 2009) showed reduced quenching of EGFP in shPAT2 and PAT2-HA adipocytes upon amino acid starvation for one hour, when expressed as EGFP/mCherry ratio (**Fig. 3F** **and Fig. S4E**).

**Figure 3:**
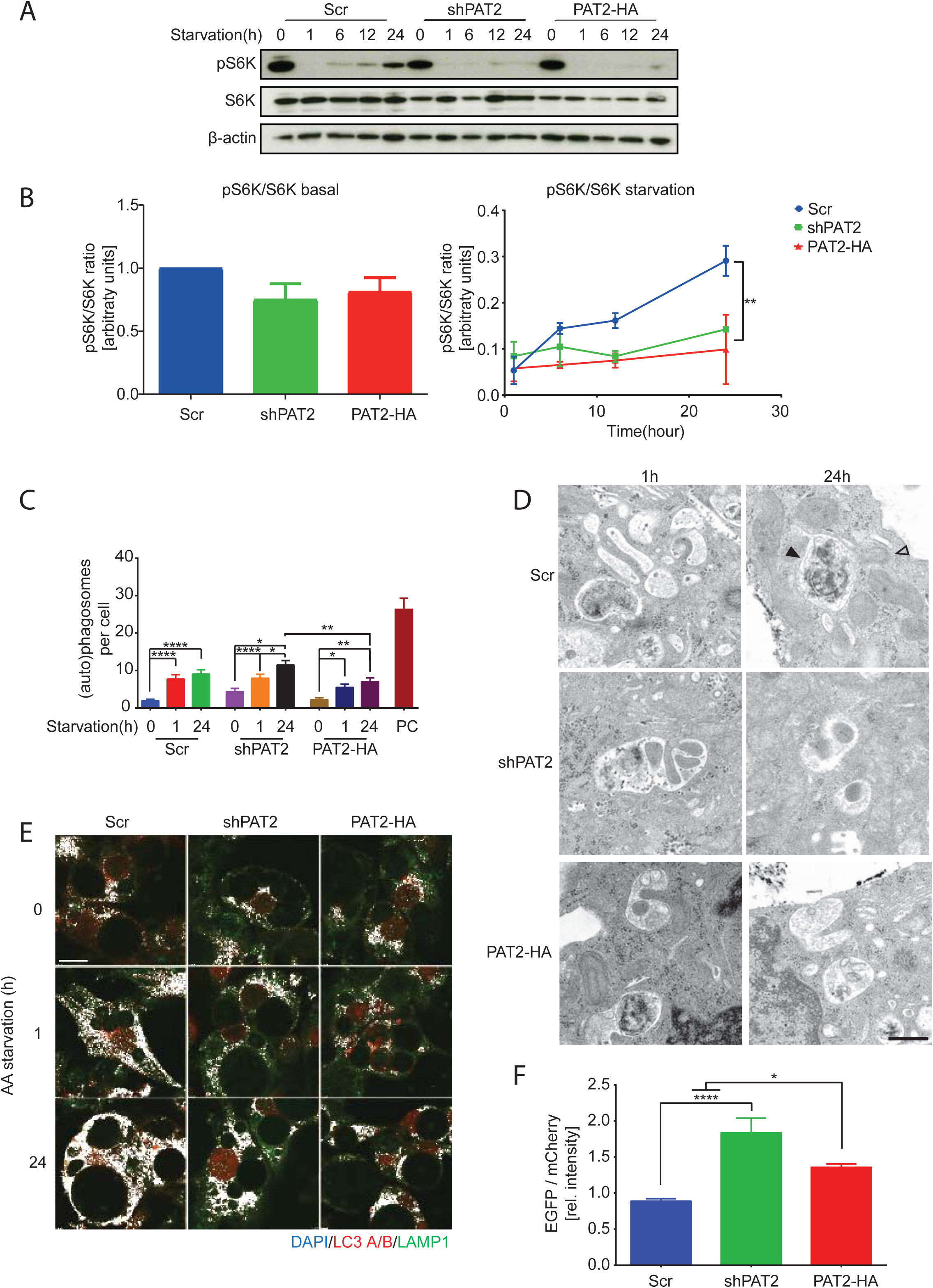
Knockdown and overexpression of PAT2 impair autophagy in brown adipocytes. **(A)** Western blot for S6K phosphorylation in Scr, shPAT2 and PAT2-HA brown adipocytes upon amino acid and serum depletion for 1,6,12 or 24 hours. **(B)** Quantification of relative S6K phosphorylation at baseline (left panel) or during starvation (right panel, normalized to total S6K and fold Scr; n=3; ** p<0.01; RM two-way ANOVA with Tukeys’ post hoc test; error bars show SEM. **(C)** Quantification of autophagosome number per cell in Scr, shPAT2 and PAT2-HA adipocytes in regular culture conditions and upon one or 24 hours amino acid starvation (n=30 cells per condition; positive control (PC) was excluded from statistic, RM two-way ANOVA with Tukeys’ post hoc test; **** p<0.0001, ** p<0.01, * p<0.05,error bars show SEM). **(D)** Representative electron microscopy images for Scr, shPAT2 and PAT2-HA cells upon amino acid starvation for one and 24 hours. Black arrow indicates autophagosomes, open arrow indicates lysosome, scale bar shows 500 nm. **(E)** Colocalization of LAMP1 and LC3 A/B immunostaining in Scr, shPAT2 and PAT2-HA adipocytes upon 0, 1 and 24 hours amino acid starvation. Colocalized pixels are shown in white. Scale bar shows 10 µm. **(F)** Quantification of EGFP quenching, in cells transiently transfected with mCherry-EGFP-LC3, upon one hour amino acid starvation (n = 28 - 33 cells per condition, **** p<0.0001, * p<0.05, one-way ANOVA, Tukeys’ post hoc test; error bars show SEM).

These data suggest that PAT2 overexpression or knockdown could impair lysosomal acidification or autophagosome and lysosome fusion. Abnormal lysosomal pH impairs autophagosome function (Kissing et al., 2015), and could explain the observed phenotypes of shPAT2 and PAT2-HA adipocytes. Assessment of intracellular pH, using a pH sensitive dye in brown adipocytes, transiently transfected with EGFP-LC3 to visualize autophagolysosomes, revealed decreased lysosomal pH in control cells following amino acid starvation, whereas this was strongly blunted in shPAT2 cells (**Fig. 4A** **and Fig. S5A**). In contrast, PAT2-HA adipocytes showed decreased lysosomal pH already in regular culture conditions (**Fig. 4A** **and Fig. S5A**). Thus, albeit both knockdown and overexpression of PAT2 both resulted in impaired starvation dependent reactivation of S6K, as a surrogate for amino acid release from the autophagolysosome, the underlying mechanism appears opposed. Loss of PAT2 increases lysosomal pH, whereas overexpression of PAT2-HA results in hyperacidification of the lysosome. Albeit, PAT2 in itself is able to transport protons, lysosomal acidification is thought to be predominantly driven by the V-ATPase (Mindell, 2012). Indeed, vATPase inhibition using bafilomycin A1 confirmed the dependency of starvation induced lysosomal acidification on vATPase in our cell lines (**Fig. S5B**).

**Figure 4:**
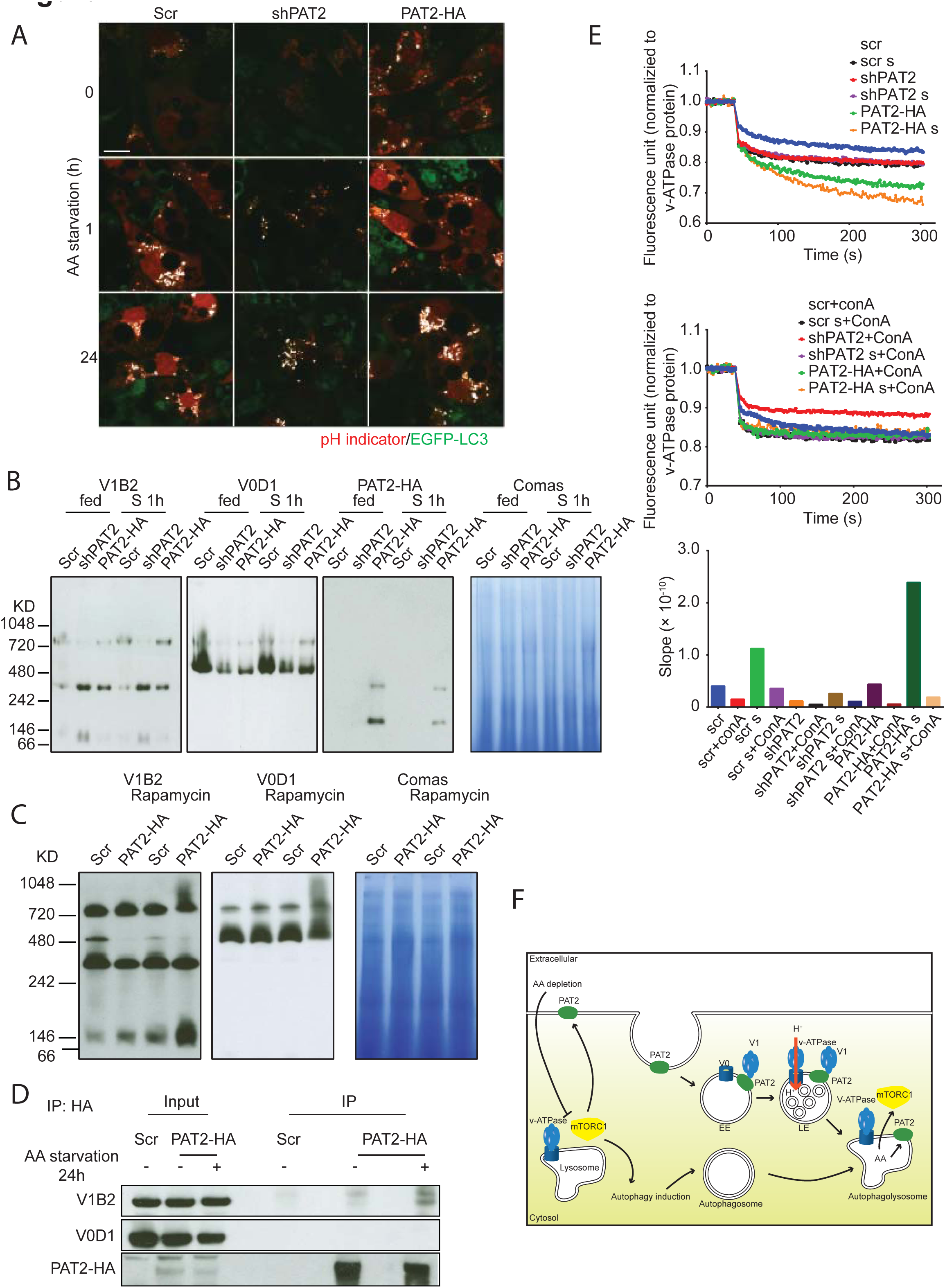
PAT2 regulates assembly and proton pumping efficiency of the lysosomal vATPase. **(A)** Fluorescence images of Scr, shPAT2 and PAT2-HA brown adipocytes transiently transfected with EGFP-LC3 and stained with an intracellular pH indicator in control medium or amino acid and serum free medium for 1 or 24 hours. Colocalized pixels are shown in white. Scale bar shows 15µm. **(B)** BN-PAGE of whole cell lysates from Scr, shPAT2 and PAT2-HA brown adipocytes in control or amino acid free DMEM (1 hour). **(C)** BN-PAGE for v-ATPase assembly in Scr and PAT2-HA adipocytes following one hour rampamycin (10µM) treatment. **(D)** Co-immunoprecipitation of PAT2-HA with components of the lysosomal vATPase in differentiated brown adipocytes. **(E)** Lysosomal and endosomal FITC-dextran quenching in Scr, shPAT2 and PAT2-HA brown adipocytes in control and amino acid free DMEM (1 hour). Acute treatment with 1 µM Concanamycin A was used as negative control. Lysosomal acidification rates shown as slopes were calculated from regression analysis of the samples and curves. **(F)** The model for PAT2 mediated v-ATPase assembly in response to amino acid depletion in brown adipocytes.

Western blots did not show differences in protein levels of V1B2 between Scr, shPAT2 and PAT2-HA adipocytes (**Fig. S5C**). However, regulation of vATPase assembly is the main mechanism regulating vATPase activity (McGuire, Stransky et al., 2017). Using blue native PAGEs we detected the fully assembled vATPase at >720kDa, as determined by an overlapping signal of V1B2 and V0D1 (**Fig. 4B**). Both control and PAT2-HA cells increased the amount of fully assembled vATPase upon amino acid starvation, whereas shPAT2 cells had strongly decreased amounts of vATPase both at baseline and upon amino acid starvation. Thus, the observed increase in lysosomal pH in shPAT2 adipocytes appears as the consequence of impaired assembly of the full size vATPase. Membrane and cytosol fractionation independently confirmed reduced vATPase assembly upon amino acid depletion in shPAT2 adipocytes (**Fig. S5D**). Importantly, the role of PAT2 to regulate vATPase assembly is independent of mTORC1 activity, as rapamycin treatment alone was insufficient to trigger vATPase assembly (**Fig. 4C**). Mechanistically we show using co-immunoprecipitations that PAT2-HA interacts with the V1B2 subunit of the v-ATPase V1 domain, but not V0D1, a V0 subunit, in amino acid starved adipocytes (**Fig. 4D**). This suggests a mechanism whereby PAT2, upon translocation from the plasma membrane via the endosome to the lysosome, facilitates the assembly of the full length vATPase by recruiting the V1 domain to the lysosomal surface, where it interacts with the V0 subunit. Interestingly, full size vATPase was only marginally increased in PAT2-HA cells, especially in fed conditions when compared to control cells. Thus, assembly alone cannot fully explain the hyperacidification observed upon PAT2 overexpression. Therefore, we tested if PAT2, in addition to vATPase assembly, can also regulate vATPase proton pumping efficiency. Measurements of pH dependent quenching of FITC-dextran (Stransky & Forgac, 2015) normalized to intact vATPase (**Fig. S5E**) showed enhanced proton pumping in response to amino acid starvation in control and PAT2-HA cells with greatly increased pumping efficiency in PAT2-HA cells (**Fig. 4E**), indicating that decreased lysosomal pH in PAT2 overexpressing brown adipocytes is the result of increased vATPase pumping efficiency rather than increased assembly.

In summary we identify the amino acid transporter PAT2 to promote lysosomal acidification upon reductions in extracellular amino acid availability in brown adipocytes. In response to amino acid depletion induced mTORC1 inhibition, PAT2 translocates from the plasma membrane to the lysosome, where it facilitates the assembly of the vATPase promoting lysosomal acidification, which is essential for the induction of autophagy (**Fig. 4F**). Thus, the very high expression of PAT2 in brown adipose tissue and the here described functions suggest that BAT has a very high sensitivity towards changes in extracellular amino acid levels to regulate its thermogenic function, providing previously unrecognized opportunities for the pharmacological modulation of BAT activity *in vivo*.

## Materials and Methods

### Cell culture

For all experiments a previously established murine brown preadipocyte cell line, derived from an 8 week old C57Bl/6 mouse was used, cultured and differentiated as previously described(Pramme-Steinwachs, Jastroch et al., 2017). To establish a PAT2 knockdown cell line, a shPAT2 (targeting sequence:

CCGGCAGACTGAACAAGCCTTTCATCTCGAGATGAAAGGCTTGTTCAGTCTGTTTTTG) and its scrambled control shRNA, cloned into a pLKO.1-puro vector were purchased from Sigma Aldrich. PAT2 cDNA containing a HA-tag directly in front of the stop codon was cloned into the pCDH-CMV-puro plasmid to generate the PAT2-HA overexpression cell line. All plasmids were packed in lentiviruses, concentrated using PEG-it (SystemBio) and preadipocytes were infected in presence of 9 μg/ml polybrene. Cells were cultured in medium containing DMEM, 10% fetal bovine serum, 1% penicillin-streptomycin and 2.5 μg/ml puromycin.

#### Adipocyte differentiation and amino acid starvation

Preadipocytes were grown to 100 % confluence and the differentiation was induced with 0.5 mM IMBX, 5 µM dexamethasone, 0.125 mM indomethacine (Santa Cruz Biotechnology), 1 nM triiodothyronine (T3, Merck Millipore), 100 nM insulin and 1 nM rosiglitazone. After two days, the medium was changed to medium containing only 100 nM insulin and 1 nM T3. The medium was changed every 2 days until day 8. For amino acid starvation, cells were washed twice with PBS and cultured with amino acid free DMEM (GENAXXON bioscience) containing 1% penicillin-streptomycin and dialyzed FBS (Thermo Scientific) if indicated. Amino acid restimulation was performed by adding MEM Amino Acids (50x) solution (Sigma Aldrich).

#### Transient transfection

30 µl DMEM, 20 µl Polyfect (Qiagen) and 1 µg plasmid were incubated for 5 minutes and the transfection mix was dropped to cover all cells without medium. After 4 h incubation cell culture medium was added.

#### Proliferation assay

2000 cells per well were plated in 96 well plates in 500 μl medium and grown for 1-4 days. 50 μl Cell Counting Kit – 8 solution (Sigma Aldrich) was added to each well and cells were incubated for 1 h in the cell culture incubator and absorptions at 450 nm was measured.

### Oil Red O staining

Cells were fixed with 10 % formalin in PBS for 10 min, washed with PBS twice and incubated with 60% isopropanol for 5 min. Cells were incubated in 21% Oil Red O (Sima Aldrich) in 60% isopropanol for 10 min followed by 4 time washing with distilled water. Images were taken using an EVOS XL Core Cell Imaging System (Thermo Fisher Scientific). Oil Red O was extracted by 100% isopropanol and quantified at 505 nm.

### Western blot and co-immunoprecipitation

Cells or tissues were lysed in ice-cold RIPA buffer [50mM Tris (pH=7,4), 150mM NaCl, 1mM EDTA, 1% Triton X100] containing 1% protease and phosphatase inhibitor cocktails (Sigma Aldrich) on ice. Protein concentrations were measured by BCA assay (Thermo Fisher Scientific). Protein samples were mixed with sample buffer (Thermo Fisher Scientific) and incubated at 70*°*C for 10 min. Proteins were transferred to 0.45 μm PVDF membranes (Merck Millipore), and blocked with 5% skim milk in TBS-0.1% Tween20 (TBST) for at least one hour. Primary and HRP conjugated secondary antibodies (Table S1) were diluted in 5% BSA in TBST. Amersham Hyperfilm ECL (GE) and HRP substrate ECL (Merck Millipore) were used to detect signals. Band intensities were quantified by ImageJ.

#### Co-immunoprecipitation

Cells were lysed in Pierce IP Lysis Buffer (25 mM Tris HCl pH 7.4, 150 mM NaCl, 1% NP-40, 1 mM EDTA, 5% glycerol) (Thermo Fisher Scientific) containing 1% protease and phosphatase inhibitors on ice and protein concentration was measured by BCA. 1 mg of protein lysate was incubated with 1μg anti HA-antibody (Roche) overnight at 4*°*C. 10 μl Dynabeads protein G (Santa Cruz) were added to the lysate for 1 h at 4*°*C. The beads were precipitated by centrifugation at 1000 × g at 4*°*C for 3 min. Lysis buffer was used to wash beads 3 times and proteins were eluted with NuPAGE™ LDS Sample Buffer (2X) with 5% β-mercaptoethanol at 70*°*C for 5min and analyzed by western blot. For MS sample preparation, beads were additionally washed twice with buffer containing 25 mM TrisHCl (pH 7.4), 150 mM NaCl, 1 mM EDTA, 5% glycerol, 1% protease and phosphatase inhibitors before elution of the proteins with NuPAGE™ LDS Sample Buffer (2X) with 5% β-mercaptoethanol at 70*°*C for 5min.

### Blue native-PAGE

The NativePAGE Novex Bis-Tris gel system (Thermo Fisher Scientific) was used according to the manufacturer’s instruction using 1% digitonin and NativePAGE 3-12% Bis-Tris gels. Gels were soaked in 0.1% SDS in TBST for 10 min before transfer to 0.45 µm PVDF membranes unsing a Bio Rad wet tank blotting system with 0.01% SDS in Tris glycine transfer buffer containing 10% methanol. The membrane was incubated in 8% acetic acid in TBST for 15 min and subsequently washed with double distilled water. The remaining Coomassie G250 dye was removed with 100% methanol and the membrane used for western blot.

### Subcellular fractionations

Subcellular fractionation was performed as previously described(Stransky & Forgac, 2015). In brief, cells were homogenized in fractionation buffer (250 mM sucrose, 1 mM EDTA and 10 mM HEPES pH 7.4) containing protease and phosphatase inhibitors using a Potter-Elvehjehm grinder on ice. Homogenates were centrifuged at 500 × g for 10 min at 4°C and the supernatant at 100000 × g for 30 min at 4°C to pellet membranes. The cytosol fraction in the supernatant was concentrated using 10K Polyethersulfone (PES) membranes (VWR) according to the manufacturer’s instructions. The membrane pellet was washed with fractionation buffer. 0.1% SDS was added to the cytosol and membrane fractions and analyzed by western blot.

### Fluorescence stainings and imaging

Cells were cultured on chamber slides (Thermo Fisher Scientific), fixed with 4% PFA (Sigma Aldrich) or methanol for 10 min. Tissues were fixed with 4% PFA for 1 h prior to vibratome (Leica) sectioning at 100 μm. Cells or tissue sections were washed with PBS and 3% BSA and 0.3%Tween 20 in PBS were used for blocking and permeabilisation for 1 h. Samples were incubated with primary antibodies overnight and Alexa conjugated secondary antibodies (see Table S1) for one hour. DAPI diluted in PBS (1 : 5000) was added to the cells after the secondary antibody for 5 min. Cells and tissue sections were mounted with mounting medium (Dako) and images acquired using a Leica TCS SP5 confocal microscope. Image quantification and co-localization analysis were performed using ImageJ.

#### EGFP quenching

pBABE-puro mCherry-EGFP-LC3B deposited by Jayanta Debnath lab was obtained from Addgene (# 22418)(N’Diaye et al., 2009). The plasmid was transiently transfected into adipocytes cultured in live cell imaging chamber slides (ibidi). Following the amino acid starvation for one hour, the medium was changed to live cell imaging solution (Thermo Fisher Scientific) and images were acquired by confocal microscopy maintaining 5% CO2 and 37°C during imaging. Relative intensities of mCherry and EGFP were quantified by ImageJ software.

#### Intracellular pH measurements

pEGFP-LC3 (human) deposited by Toren Finkel lab was obtained from Addgene (# 24920)(Lee, Cao et al., 2008). The plasmid was transiently transfected into adipocytes. pHrodo Red AM (Thermo Fisher Scientific) was used to assess intracellular pH following the manufacturer’s instruction and imaged as described above. Images were analyzed for co-localization of red and green pixels by ImageJ software.

#### In vitro quenching test

The protocol was modified from the published method(Stransky & Forgac, 2015) as described. 2.2 mg/ml FITC-Dextran 70000 (Sigma Aldrich) in culture medium was added to adipocytes overnight. The medium was replaced with culture medium or amino acid free DMEM for 1 h. The adipocytes were homogenized in 125 mM KCl, 1 mM EDTA, 50 mM sucrose, 20 mM HEPES pH 7.4, 1% phosphatase inhibitor cocktails and protease inhibitor cocktail using a Potter-Elvehjehm grinder on ice. Big particles were removed by centrifugation at 2000 × g for 10 min at 4°C. The FITC-Dextran loaded vesicles were pelleted by centrifugation at 16100 × g for 15 min at 4°C and the pellet resuspended in homogenization buffer. Protein concentration was measured using a BCA kit. Particles corresponding to 4 µg protein were added with or without 1 µM concanamycin A (Santa Cruz) to flat glass bottom plates to measure FITC fluorescence at 488 nm at 37°C. Fluorescence intensity was recorded every 2 s for 30 cycles and for additional 120 cycles after addition of 10 mM ATP and 20 mM MgCl_2_. Data were normalized to relative vATPase quantity as assessed by BN-PAGE of the same samples. All intensities were normalized to the baseline of Scr adipocytes samples cultured in regular culture medium. Data transformation was performed according to the Stern-Volmer equation(Stern, 1919) and slopes subsequently calculated.

### Semiquantitative Realtime PCR

RNA was extracted from cells and tissues using the RNeasy kit (Qiagen) following the manufacturer’s recommendations. RNA concentrations were measured using a Nano Drop 2000 (Thermo Fisher Scientific) and cDNA was synthesized using 500-1000 ng RNA and the High-capacity cDNA reverse transcription kit (ABI) according to the manufacturer’s instruction. SYBR green (Bio Rad) based real time PCR was performed by using a CFX384 Touch™ Real-Time PCR Detection System (Bio Rad) with the program 95°C 30 s, (95°C 5 s, 60°C 30 s) × 40 cycles, 95°C 10 s. Primers sequences: pat2 forward (f) GTGCCAAGAAGCTGCAGAG, reverse (r) TGTTGCCTTTGACCAGATGA; tbp f ACCCTTCACCAATGACTCCTATG, r TGACTGCAGCAAATCGCTTGG; pparγ f TCCTATTGACCCAGAAAGCGA, r TGGCATCTCTGTGTCAACCA; lc3b f AGAGTGGAAGATGTCCGGCT, r TCTCCCCCTTGTATCGCTCT.

### Electron microscopy

Preadipocytes were plated on collagen I coated coverslips and differentiated. For electron microscopy, the cells were fixed using 4% paraformaldehyde (Serva, Heidelberg, Germany) and 2% glutaraldehyde (Serva) in PBS followed by staining with 0.5% osmium tetroxide (EMS, Hatfield, PA, USA). After thorough rinsing in PBS, the sections were dehydrated in graded alcohol and further stained with 1% uranyl acetate (Merck, Darmstadt, Germany) in 70% alcohol. After final dehydration the samples were transferred in propylene oxide (Sigma Aldrich, Steinheim, Germany) and incubated in Durcupan (Sigma Aldrich). After polymerization at 56°C for 48h, the cell culture insert was removed and the blocks of resin containing the embedded cells were trimmed and finally cut using an ultra-microtome (Leica Microsystems, Wetzlar, Germany). Ultra-thin sections with an average thickness of 55nm were transferred on formvar-coated copper grids and stained with lead citrate. Analysis was performed using a Zeiss SIGMA electron microscope (Zeiss NTS, Oberkochen, Germany) equipped with a STEM detector and ATLAS software.

### Statistical analysis

GraphPad PRISM 6 was used for statistical analysis. Error bars, P values, sample size and statistical tests are detailed in the respective figure legends.

## Acknowledgements

This work was supported by iMed the initiative for personalized medicine of the Helmholtz Association and funds from the German research foundation (DFG) as well as from the project Aging and Metabolic Programming (AMPro). JW was supported by the China Scholar Council (CSC). **Author contributions:** JW and SU designed and conducted the experiments and wrote the manuscript. MK conducted the electron microcopy experiments. SMH contributed to study design and data analysis. **Competing financial interests:** The authors declare no conflict of interest.

---

**Figurs S1.**
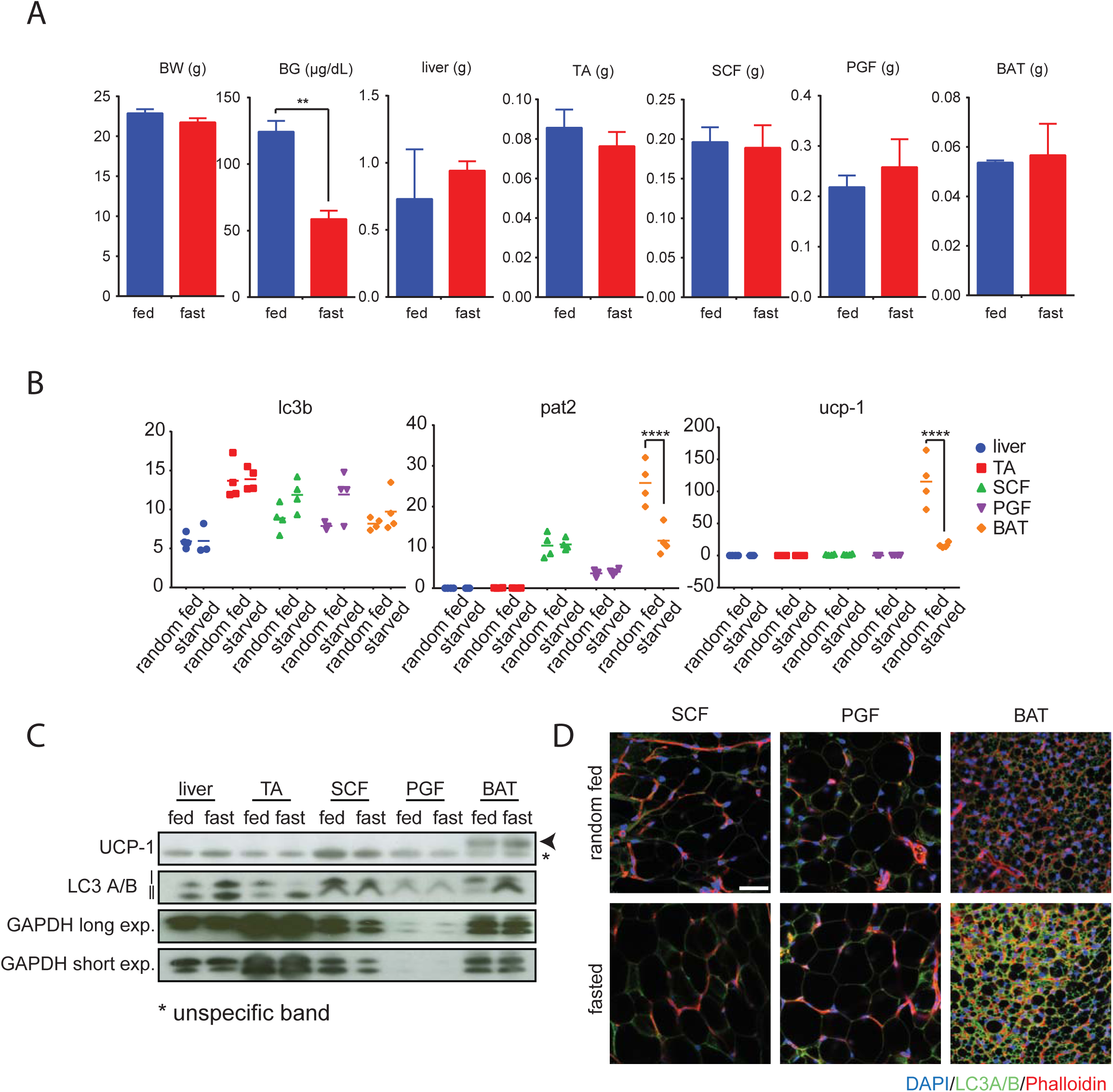
**(A)** Body weights, blood glucose and weights of liver, tibialis anterior (TA), subcutaneous fat (SCF), perigonadal fat (PGF) and brown adipose tissue (BAT)) from ad libitum fed and overnight fasted mice (n=3). ** p<0.01; unpaired t-test; Error bars show SEM. **(B)** Semiquantitative PCR of LC3B, PAT2 and UCP-1 expression in liver, TA, SC, PG and BAT from ad libitum fed and overnight fasted mice. n=3-4; **** p<0.0001; RM two-way ANOVA with Tukeys’ post hoc test. **(C)** Western blots from ad libitum fed and overnight fasted mice (liver, tibialis anterior (TA), subcutaneous fat (SCF), perigonadal fat (PGF) and brown adipose tissue (BAT)). **(D)** Immunofluorescence stainings of LC3A/B and DAPI for adipose tissues from ad libitum fed and overnight fasted mice. Scale bar shows 40 μm.

**Figure S2.**
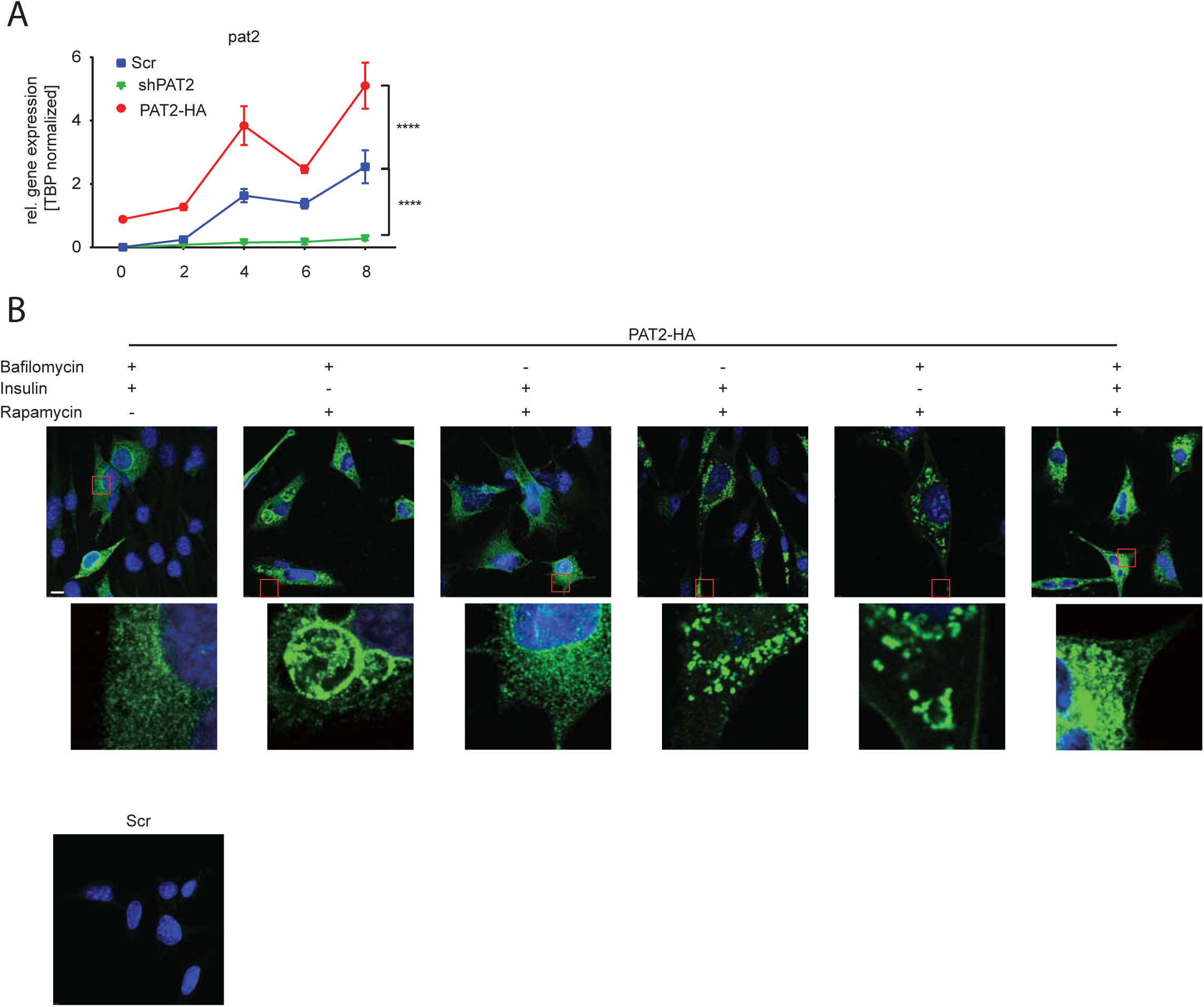
**(A)** Semiquantitative PCR of PAT2 for Scr, shPAT2 and PAT2-HA brown adipocytes at indicated time points during differentiation (n=3). **** p<0.0001; RM Two-way ANOVA with Tukeys’ posthoc test. **(B)** Immunostaining for PAT2-HA in preadipocytes treated with Bafilomycin A1 (100 nM), insulin (100 nM), Rapamycin (10 µM) combinations. Scr preadipocytes were used as negative control. Scale bar shows 10 µm.

**Figure S3.**
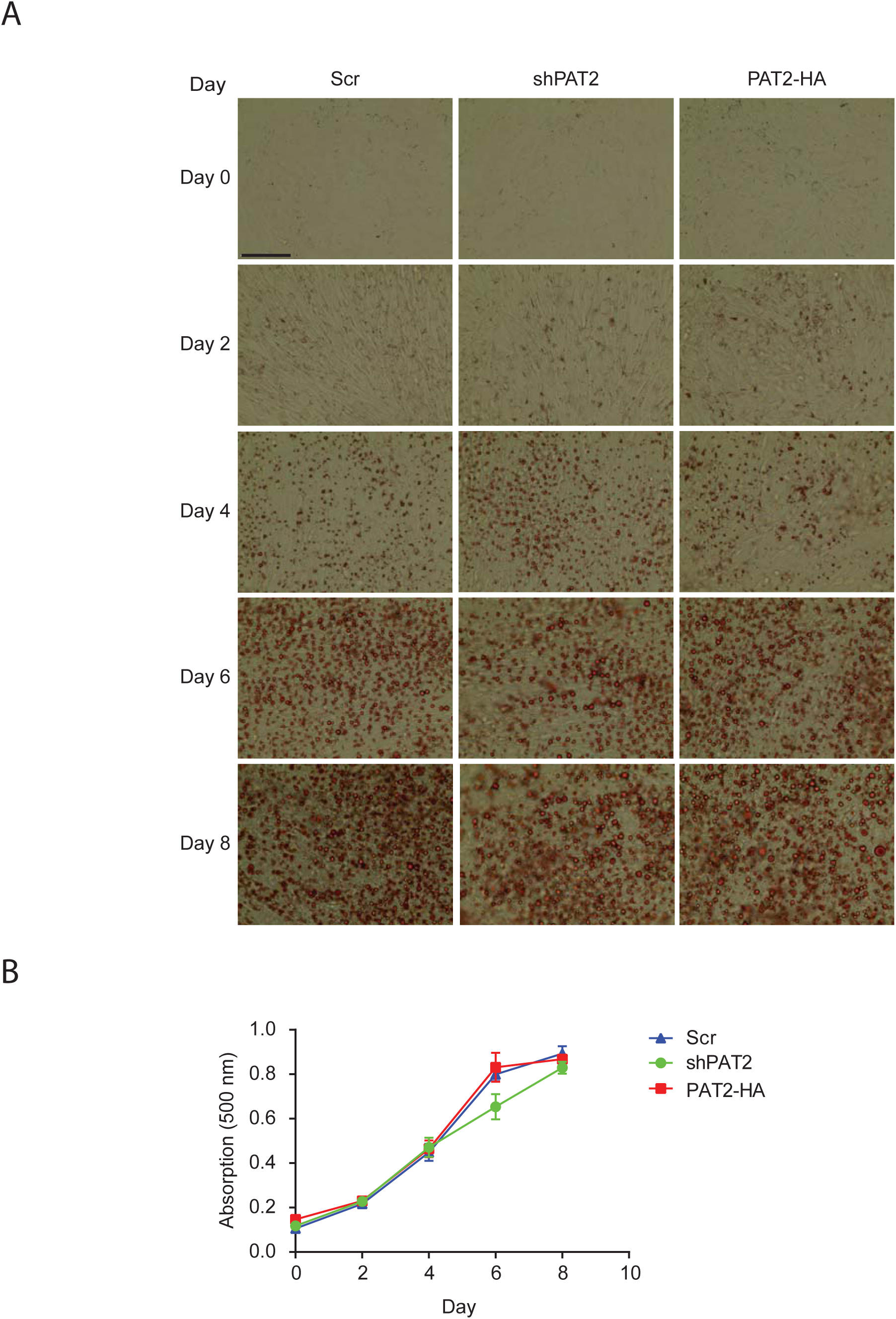
**(A)** Lipid accumulation during brown adipocyte differentiation visualized by Oil Red O staining. Scale bar shows 100 µm. **(B)** Quantification of Oil Red O stain from (A) (n=3).

**Figure S4.**
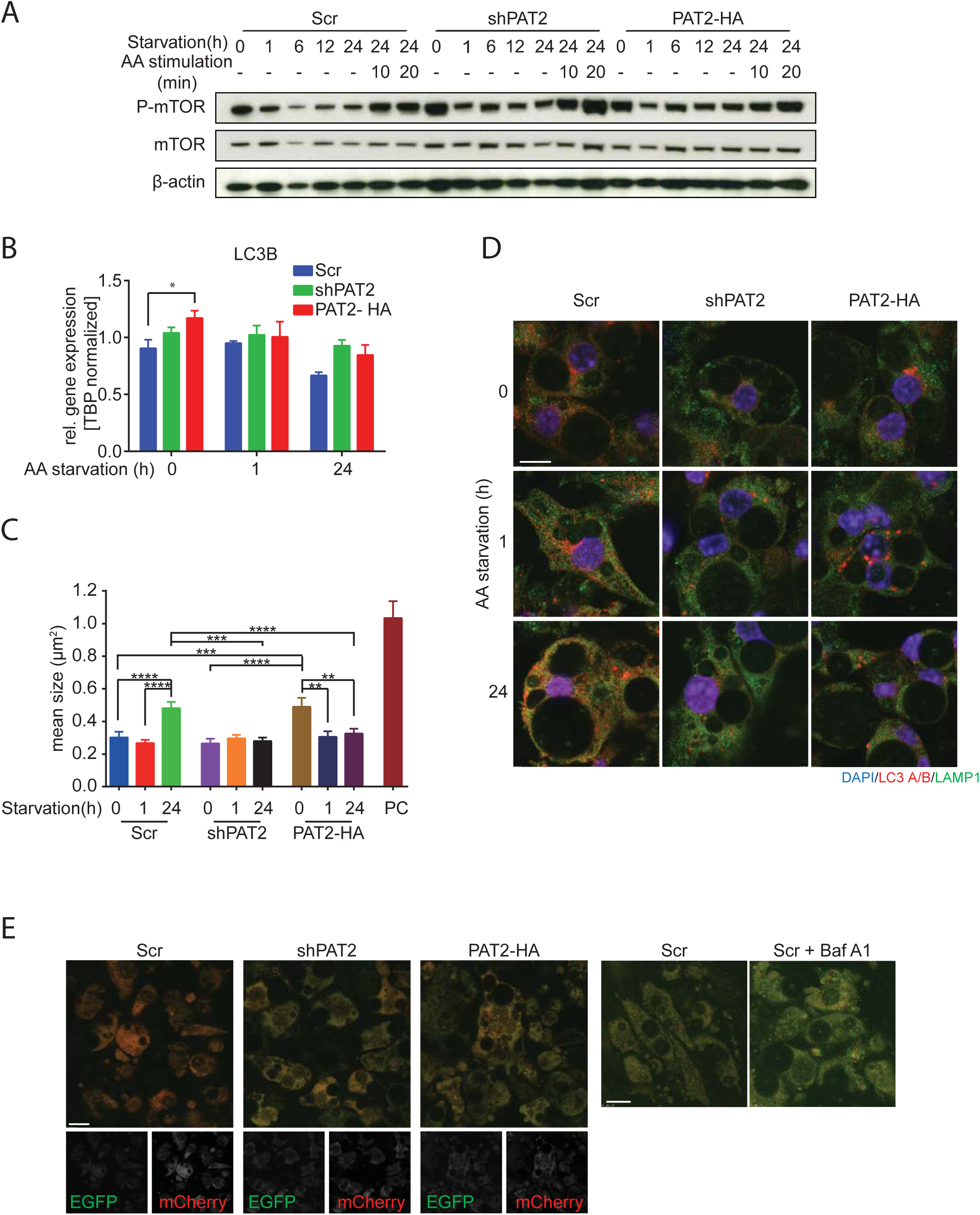
**(A)** Western blot for phosphorylated mTOR (Ser2448), mTOR and β-actin in Scr, shPAT2 and PAT2-HA upon amino acids and serum depletion for the indicated times, followed by restimulation with amino acids for 10 or 20 min after 24 hours starvation. **(B)** LC3B expression during amino acid and serum depletion (n=3; * p<0.05; RM two-way ANOVA with Tukeys’ post hoc test; error bars show SEM). **(C)** Quantification of average autophagosome size (n=74-161) in Scr, shPAT2 and PAT2-HA adipocytes in regular culture conditions and upon one or 24 hours amino acid starvation (Positive control (PC) was excluded from statistic, RM two-way ANOVA with Tukeys’ post hoc test; **** p<0.0001, *** p<0.001, ** p<0.01, error bars show SEM). **(D)** Original images of Figure 3D. **(E)** Visualization of autophagosome and lysosome fusion by assessing quenching of EGFP in cells transiently transfected with mCherry-EGFP-LC3 following 24 hours amino acid starvation. Control cells were additionally treated with Bafilomycin A1 (Baf A1, 100nM). Scale in left panel shows 15 µm and in right panel shows 10 µm.

**Figure S5.**
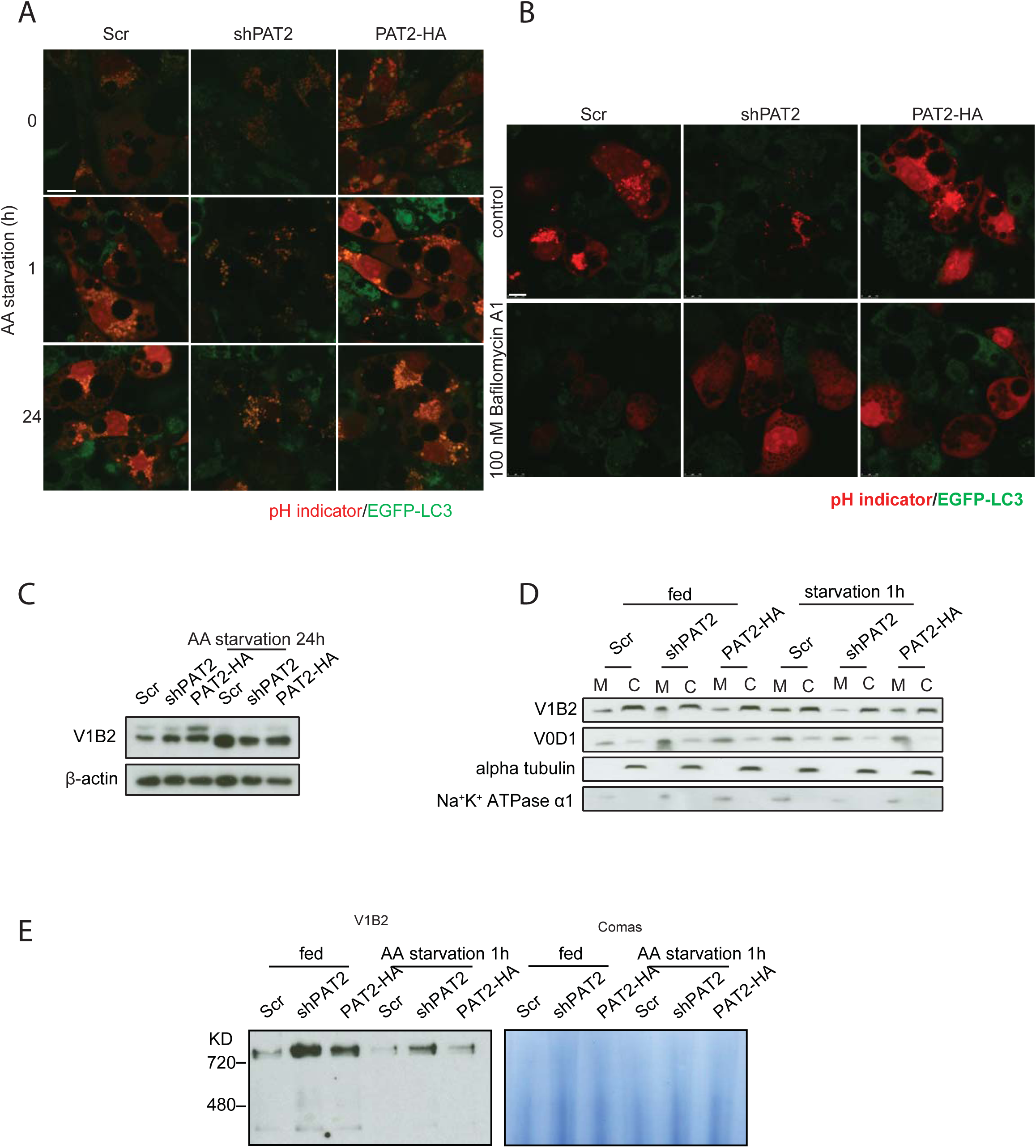
**(A)** Representative images for Figure 4A. **(B)** Fluorescence images of cells following 24 h amino acid and serum starvation in presence or absence of the v-ATPase inhibitor Bafilomycin A1 (100nM). Scale bar shows 7.5 µm. **(C)** WB for V1B2 in Scr, shPAT2 and PAT2-HA brown adipocytes cultured in normal medium or amino acids free DMEM for 24 hours. **(D)** Western blot for the cytosol (C) and membrane (M) fractions from control and amino acid starved (1 hour) Scr, shPAT2 and PAT2-HA adipocytes. **(E)** Western blot for V1B2 following a BN-PAGE of v-ATPase for normalization in Figure 4E.

**Supplemental Table 1.**

|  |  |  |
| --- | --- | --- |
| HA-Tag (6E2) Mouse mAb (HRP Conjugate) | WB 1 : 2000 | Cell Signaling Technology |
| HA-tag Rat (12CA5) | IP 1 : 200 | Roche |
| PPARγ (C26H12) Rabbit mAb | WB 1 : 1000 | Cell Signaling Technology |
| UCP-1 Rabbit | WB 1 : 5000 | Gift from Prof. Martin Jastroch |
| Anti-LAMP1 antibody Rabbit polyclonal | WB 1 : 1000<br>IF 1 : 100 | Abcam |
| EEA1 (C45B10) Rabbit mAb | IF 1 : 100 | Cell Signaling Technology |
| Golgin-97 (D8P2K) Rabbit mAb | IF 1 : 100 | Cell Signaling Technology |
| mTOR (7C10) Rabbit mAb | WB 1 : 1000<br>IF 1 : 100 | Cell Signaling Technology |
| Phospho-mTOR (Ser2448) (D9C2) XP Rabbit mAb | WB 1 : 1000 | Cell Signaling Technology |
| β-Actin (C4) HRP | WB 1 : 5000 | Santa-Cruz |
| Phospho-p70 S6 Kinase (Thr389) (108D2) Rabbit mAb | WB 1 : 1000 | Cell Signaling Technology |
| p70 S6 Kinase (49D7) Rabbit mAb | WB 1 : 1000 | Cell Signaling Technology |
| LC3A/B (D3U4C) XP Rabbit mAb | WB 1 : 1000 | Cell Signaling Technology |
| Anti-LAMP1 antibody [H4A3] Mouse monoclonal | IF 1 : 10 | Abcam |
| RagC (D31G9) XP Rabbit mAb | WB 1 : 1000 | Cell Signaling Technology |
| ATP6V1B2 (D3O7Q) Rabbit mAb | WB 1 : 1000 | Cell Signaling Technology |
| Recombinant Anti-ATP6V0D1 /P39 antibody [EPR18320-38] Rabbit monoclonal | WB 1 : 2000 | Abcam |
| GAPDH Antibody (D-6) HRP | WB 1 : 5000 | Santa-Cruz |
| OxPhos Rodent WB Antibody Cocktail | WB 1 : 1000 | Thermo Fisher Scientific |
| Anti-rabbit IgG, HRP-linked Antibody | WB 1 : 5000 | Cell Signaling Technology |
| goat anti-mouse IgG-HRP | WB 1 : 5000 | Santa-Cruz |
| Alexa Fluor 488 AffiniPure Donkey Anti-Rat IgG (H+L) | IF 1 : 500 | Jackson ImmunoResearch |
| Alexa Fluor 488 AffiniPure Rat Anti-Mouse IgG (H+L) | IF 1 : 500 | Jackson ImmunoResearch |
| Alexa Fluor 647 Goat<br>Anti-Rabbit (H+L) | IF 1 : 500 | Thermo Fisher<br>Scientific |
| Alexa Fluor 594<br>AffiniPure Goat Anti-Rabbit IgG (H+L) | IF 1 : 500 | Jackson<br>ImmunoResearch |

## References

1. Abu-Remaileh M, Wyant GA, Kim C, Laqtom NN, Abbasi M, Chan SH, Freinkman E, Sabatini DM (2017) Lysosomal metabolomics reveals V-ATPase- and mTOR-dependent regulation of amino acid efflux from lysosomes. Science 358: 807–813

2. Ben-Sahra I, Manning BD (2017) mTORC1 signaling and the metabolic control of cell growth. Curr Opin Cell Biol 45: 72–82

3. Boll M, Foltz M, Rubio-Aliaga I, Kottra G, Daniel H (2002) Functional characterization of two novel mammalian electrogenic proton-dependent amino acid cotransporters. J Biol Chem 277: 22966–73

4. Broer S, Broer A (2017) Amino acid homeostasis and signalling in mammalian cells and organisms. Biochem J 474: 1935–1963

5. Condon KJ, Sabatini DM (2019) Nutrient regulation of mTORC1 at a glance. J Cell Sci 132

6. Dikic I, Elazar Z (2018) Mechanism and medical implications of mammalian autophagy. Nat Rev Mol Cell Biol 19: 349–364

7. Ferhat M, Funai K, Boudina S (2018) Autophagy in Adipose Tissue Physiology and Pathophysiology. Antioxid Redox Signal

8. Foltz M, Oechsler C, Boll M, Kottra G, Daniel H (2004) Substrate specificity and transport mode of the proton-dependent amino acid transporter mPAT2. Eur J Biochem 271: 3340–7

9. Galluzzi L, Pietrocola F, Levine B, Kroemer G (2014) Metabolic control of autophagy. Cell 159: 1263–76

10. Goberdhan DC, Wilson C, Harris AL (2016) Amino Acid Sensing by mTORC1: Intracellular Transporters Mark the Spot. Cell Metab 23: 580–9

11. Hankir MK, Cowley MA, Fenske WK (2016) A BAT-Centric Approach to the Treatment of Diabetes: Turn on the Brain. Cell Metab 24: 31–40

12. Hankir MK, Klingenspor M (2018) Brown adipocyte glucose metabolism: a heated subject. EMBO Rep 19

13. Heeren J, Scheja L (2018) Brown adipose tissue and lipid metabolism. Curr Opin Lipidol 29: 180–185

14. Hoeke G, Kooijman S, Boon MR, Rensen PC, Berbee JF (2016) Role of Brown Fat in Lipoprotein Metabolism and Atherosclerosis. Circ Res 118: 173–82

15. Kennedy DJ, Gatfield KM, Winpenny JP, Ganapathy V, Thwaites DT (2005) Substrate specificity and functional characterisation of the H+/amino acid transporter rat PAT2 (Slc36a2). British journal of pharmacology 144: 28–41

16. Kissing S, Hermsen C, Repnik U, Nesset CK, von Bargen K, Griffiths G, Ichihara A, Lee BS, Schwake M, De Brabander J, Haas A, Saftig P (2015) Vacuolar ATPase in phagosome-lysosome fusion. J Biol Chem 290: 14166–80

17. Klepac K, Georgiadi A, Tschop M, Herzig S (2019) The role of brown and beige adipose tissue in glycaemic control. Mol Aspects Med 68: 90–100

18. Kuruvilla R (2019) Why brown fat has a lot of nerve. Nature 569: 196–197

19. Lee IH, Cao L, Mostoslavsky R, Lombard DB, Liu J, Bruns NE, Tsokos M, Alt FW, Finkel T (2008) A role for the NAD-dependent deacetylase Sirt1 in the regulation of autophagy. Proc Natl Acad Sci U S A 105: 3374–9

20. Li Y, Schnabl K, Gabler SM, Willershauser M, Reber J, Karlas A, Laurila S, Lahesmaa M, M UD, Bast-Habersbrunner A, Virtanen KA, Fromme T, Bolze F, O’Farrell LS, Alsina-Fernandez J, Coskun T, Ntziachristos V, Nuutila P, Klingenspor M (2018) Secretin-Activated Brown Fat Mediates Prandial Thermogenesis to Induce Satiation. Cell 175: 1561–1574 e12

21. Lopez-Soriano FJ, Alemany M (1989) Effect of alanine on in vitro glucose utilization by rat interscapular brown adipose tissue. Biochim Biophys Acta 1010: 338–41

22. McGuire C, Stransky L, Cotter K, Forgac M (2017) Regulation of V-ATPase activity. Front Biosci (Landmark Ed) 22: 609–622

23. Mills EL, Pierce KA, Jedrychowski MP, Garrity R, Winther S, Vidoni S, Yoneshiro T, Spinelli JB, Lu GZ, Kazak L, Banks AS, Haigis MC, Kajimura S, Murphy MP, Gygi SP, Clish CB, Chouchani ET (2018) Accumulation of succinate controls activation of adipose tissue thermogenesis. Nature 560: 102–106

24. Mindell JA (2012) Lysosomal acidification mechanisms. Annual review of physiology 74: 69–86

25. Mizushima N (2018) A brief history of autophagy from cell biology to physiology and disease. Nat Cell Biol 20: 521–527

26. N’Diaye EN, Kajihara KK, Hsieh I, Morisaki H, Debnath J, Brown EJ (2009) PLIC proteins or ubiquilins regulate autophagy-dependent cell survival during nutrient starvation. EMBO Rep 10: 173–9

27. Nedergaard J, Cannon B (2018) Brown adipose tissue as a heat-producing thermoeffector. Handbook of clinical neurology 156: 137–152

28. Oelkrug R, Polymeropoulos ET, Jastroch M (2015) Brown adipose tissue: physiological function and evolutionary significance. J Comp Physiol B 185: 587–606

29. Okla M, Kim J, Koehler K, Chung S (2017) Dietary Factors Promoting Brown and Beige Fat Development and Thermogenesis. Adv Nutr 8: 473–483

30. Perera RM, Zoncu R (2016) The Lysosome as a Regulatory Hub. Annu Rev Cell Dev Biol 32: 223–253

31. Pramme-Steinwachs I, Jastroch M, Ussar S (2017) Extracellular calcium modulates brown adipocyte differentiation and identity. Scientific reports 7: 8888

32. Ramirez AK, Lynes MD, Shamsi F, Xue R, Tseng YH, Kahn CR, Kasif S, Dreyfuss JM (2017) Integrating Extracellular Flux Measurements and Genome-Scale Modeling Reveals Differences between Brown and White Adipocytes. Cell Rep 21: 3040–3048

33. Rebsamen M, Pochini L, Stasyk T, de Araujo ME, Galluccio M, Kandasamy RK, Snijder B, Fauster A, Rudashevskaya EL, Bruckner M, Scorzoni S, Filipek PA, Huber KV, Bigenzahn JW, Heinz LX, Kraft C, Bennett KL, Indiveri C, Huber LA, Superti-Furga G (2015) SLC38A9 is a component of the lysosomal amino acid sensing machinery that controls mTORC1. Nature 519: 477–81

34. Rubio-Aliaga I, Boll M, Vogt Weisenhorn DM, Foltz M, Kottra G, Daniel H (2004) The proton/amino acid cotransporter PAT2 is expressed in neurons with a different subcellular localization than its paralog PAT1. J Biol Chem 279: 2754–60

35. Saxton RA, Sabatini DM (2017) mTOR Signaling in Growth, Metabolism, and Disease. Cell 168: 960–976

36. Schioth HB, Roshanbin S, Hagglund MG, Fredriksson R (2013) Evolutionary origin of amino acid transporter families SLC32, SLC36 and SLC38 and physiological, pathological and therapeutic aspects. Mol Aspects Med 34: 571–85

37. Stern O (1919) Uber die abklingungszeit der fluoreszenz. Phys Z 20: 183–188

38. Stransky LA, Forgac M (2015) Amino Acid Availability Modulates Vacuolar H+-ATPase Assembly. J Biol Chem 290: 27360–9

39. Suryawan A, Nguyen HV, Almonaci RD, Davis TA (2013) Abundance of amino acid transporters involved in mTORC1 activation in skeletal muscle of neonatal pigs is developmentally regulated. Amino Acids 45: 523–30

40. Thwaites DT, Anderson CM (2011) The SLC36 family of proton-coupled amino acid transporters and their potential role in drug transport. British journal of pharmacology 164: 1802–16

41. Townsend KL, Tseng YH (2014) Brown fat fuel utilization and thermogenesis. Trends Endocrinol Metab 25: 168–77

42. Ussar S, Lee KY, Dankel SN, Boucher J, Haering MF, Kleinridders A, Thomou T, Xue R, Macotela Y, Cypess AM, Tseng YH, Mellgren G, Kahn CR (2014) ASC-1, PAT2, and P2RX5 are cell surface markers for white, beige, and brown adipocytes. Sci Transl Med 6: 247ra103

43. Wanders D, Stone KP, Dille K, Simon J, Pierse A, Gettys TW (2015) Metabolic responses to dietary leucine restriction involve remodeling of adipose tissue and enhanced hepatic insulin signaling. Biofactors 41: 391–402

44. Wu X, Zhao L, Chen Z, Ji X, Qiao X, Jin Y, Liu W (2016) FLCN Maintains the Leucine Level in Lysosome to Stimulate mTORC1. PLoS One 11: e0157100

45. Wu Z, Satterfield MC, Bazer FW, Wu G (2012) Regulation of brown adipose tissue development and white fat reduction by L-arginine. Curr Opin Clin Nutr Metab Care 15: 529–38

46. Wyant GA, Abu-Remaileh M, Wolfson RL, Chen WW, Freinkman E, Danai LV, Vander Heiden MG, Sabatini DM (2017) mTORC1 Activator SLC38A9 Is Required to Efflux Essential Amino Acids from Lysosomes and Use Protein as a Nutrient. Cell 171: 642–654 e12

47. Yu L, McPhee CK, Zheng L, Mardones GA, Rong Y, Peng J, Mi N, Zhao Y, Liu Z, Wan F, Hailey DW, Oorschot V, Klumperman J, Baehrecke EH, Lenardo MJ (2010) Termination of autophagy and reformation of lysosomes regulated by mTOR. Nature 465: 942–6

48. Zhang X, Wu D, Wang C, Luo Y, Ding X, Yang X, Silva F, Arenas S, Weaver JM, Mandell M, Deretic V, Liu M (2019) Sustained activation of autophagy suppresses adipocyte maturation via a lipolysis-dependent mechanism. Autophagy: 1–15

49. Zoncu R, Bar-Peled L, Efeyan A, Wang S, Sancak Y, Sabatini DM (2011) mTORC1 senses lysosomal amino acids through an inside-out mechanism that requires the vacuolar H(+)-ATPase. Science 334: 678–83

